# TRAFfic signals: High-throughput CAR discovery in NK cells reveals novel TRAF-binding endodomains that drive enhanced persistence and cytotoxicity

**DOI:** 10.1101/2023.08.02.551530

**Authors:** Maddie D. Williams, Aye T. Chen, Matthew R. Stone, Lan Guo, Brian J. Belmont, Rebekah Turk, Nick Bogard, Nora Kearns, Mary Young, Bryce Daines, Max Darnell

## Abstract

Natural killer (NK) cells are a promising alternative therapeutic platform to CAR T cells given their favorable safety profile and potent killing ability. However, CAR NK cells suffer from limited persistence *in vivo*, which is, in part, thought to be the consequence of limited cytokine signaling. To address this challenge, we developed an innovative high-throughput screening strategy to identify CAR endodomains that could drive enhanced persistence while maintaining potent cytotoxicity. We uncovered a family of TRAF-binding endodomains that outperform benchmarks in primary NK cells along dimensions of persistence and cytotoxicity, even in low IL-2 conditions. This work highlights the importance of cell-type-specific cell therapy engineering and unlocks a wide range of high-throughput molecular engineering avenues in NK cells.

## Introduction

Chimeric antigen receptor (CAR) T cell therapies have revolutionized the landscape of hematological cancer treatment^1–4^. T cells are redirected to target and eradicate cancer in an MHC-independent manner through the introduction of CAR, translating into remarkable effectiveness against certain blood malignancies^5, 6^. Natural killer (NK) cells present several advantages over T cells, including a lower risk of graft-versus-host disease (GVHD), a common complication in donor-derived cellular therapies^7–11^. The native ability of NK cells to respond robustly to tumors without prior antigen sensitization or MHC restriction circumvents the limitations of T cell therapies in this space, namely antigen presentation and the risk of T cell anergy^12–14^. Nevertheless, to harness the full therapeutic potential of NK cells, it is essential to improve their persistence and efficacy against tumors *in vivo*^15–17^.

A critical determinant of a CAR NK cell therapy’s success is the optimal design of the CAR, a synthetic receptor that supports the recognition of cell-surface proteins in an MHC-independent manner^18, 19^. CARs can be broken down into two parts: CAR specificity is driven by the integration of an extracellular antigen-binding domain (ectodomain) and a hinge domain, while the integration of the transmembrane and one or more intracellular signaling domains (endodomains) support the induction of a signaling cascade following engagement of the receptor to its cognate antigen^20–22^. This signaling triggers a series of intracellular events leading to the release of cytolytic granules and the targeted destruction of cancer cells^23^. Despite its significance, identifying the optimal CAR endodomain configuration for potent NK cell activation remains a significant challenge^15, 24^.

Currently, CAR T cells used in a clinical setting are limited to one of two co-stimulatory endodomains, 4-1BB or CD28, along with the CD3ζ T cell activation domain. All three domains (4-1BB, CD28, and CD3ζ) have been extensively characterized in T cells, and the signaling they drive may not always translate to optimal results for NK cells due to differences in T and NK cell activation^25–30^. 4-1BB domains drive tumor necrosis factor (TNF) receptor-associated factor (TRAF) dependent signaling pathways, known to promote T cell survival and persistence. CD28 domains drive phosphatidylinositol 3-kinase (PI3K)/lymphocyte-specific protein tyrosine kinase (Lck)/growth factor receptor-bound protein 2 (Grb2) signaling pathways, known to increase T cell proliferation and cytokine production. Lastly, CD3ζ domains drive zeta-chain-associated protein kinase 70 (ZAP-70) dependent signaling pathways, known to activate T cell mediated cytotoxicity.

CAR NK cells currently used in clinical settings are engineered with CARs specifically discovered in T cell contexts. However, the different mechanisms which underlie activation, persistence, and cytotoxicity between T and NK cells motivate the urgent need to discover endodomain configurations best suited for NK cells, with the potential to enhance the potency and effectiveness of CAR NK cell therapies^31^.

Recently, endodomain screening studies were done in CAR T cells to improve their persistence and long-term efficacy, highlighting the significant role these domains play in improving CAR function^32–34^. These studies found that, despite differences, there are many convergent signaling pathways among costimulatory domains found in different immune cells. For instance, endodomains from B cell-specific TNF receptors (TACI and BAFF-R) were shown to improve CAR T cell functions^33^. We hypothesized that engineering novel CARs from a broad array of pan-immune cell receptors and signaling molecules would also yield improvements to CAR NK cell persistence and cytotoxicity.

Emerging high-throughput pooled screening methods in immunotherapy offer an opportunity to expedite the identification of novel CAR endodomains capable of substantially enhancing NK cell function^32–35^. We developed an innovative high-throughput screening strategy to identify CAR endodomains that could drive enhanced persistence while maintaining potent cytotoxicity. This technique enables simultaneous testing of a multitude of endodomain variants, accelerating the discovery process and increasing the probability of uncovering configurations that provide optimal NK cell activation. An improved understanding of how CAR endodomains influence NK cell function could help optimize current CAR NK cell therapies and pave the way for the design and engineering of next-generation CAR NK cells.

## Results

### Constructing diverse pooled CD19-CAR full-length and motif libraries

To identify CAR endodomains that could drive enhanced persistence while maintaining potent cytotoxicity, we constructed diverse pooled CD19-CAR endodomain libraries, enabling an expansive analysis of potential co-stimulatory receptor signaling. The libraries comprised a broad array of pan-immune cell receptors and signaling molecules, as well as NK-specific receptors and signaling molecules (Supplementary Tables 1 and 2).

To better appreciate how intracellular signaling influences CAR function, we built two different libraries made up of either full-length endodomains (full-length library) or smaller signaling motifs (motif library) that are known to bind to downstream signaling proteins. Each of the full-length endodomains in the full-length library contains multiple signaling motifs, and these endodomains would potentially be able to mimic their native signaling cascade upon receptor stimulation. On the other hand, each of the individual signaling motifs in the motif library may only bind to one specific downstream signaling partner^32^, and enable us to discern which binding partners or signaling pathways are utilized by the engineered CAR NK cells.

For our full-length library, we compiled a list of 36 endodomains from different signaling pathways: TRAF (4-1BB, CD27, BAFF-R, CD40, RANK, TNFRSF3, Fn14, TNFR2, CD30, TLR4, LMP1), PI3K/Fyn/Lck (CD28, DAP10, ICOS, DNAM1, 2B4, SLAMF6, SLAMF7, Ly9, CD84), Syk/ZAP-70 and their associated receptors (DAP12, FCER1G, CD3ζ, FcgRIIIa, Nkp46, NKp30, NKp44), other NK receptors (CRTAM, NKp80, CD2, IL7RA), and controls (PDCD1, OX40L, blank, NKp65 ectodomain).

For our motif library, we specifically targeted the Src Homology 2/3 (SH2/SH3)-binding domains, and TRAF-binding domains representative of both known (used in existing CAR T products) and previously unexplored signaling axes, seeking to characterize CAR NK-specific effects. We also included signaling motifs from the intracellular protein TANK, since other intracellular signaling proteins such as ZAP-70 have successfully functioned as CAR endodomains^36^.

The selected endodomains and motifs were cloned into a second-generation CAR scaffold featuring a CD19-binding single-chain variable fragment (scFv), a CD8ɑ hinge, and an NKG2D transmembrane domain (Figure 1A and 1B). Significantly, the vector used in this library did not include CD3ζ, a standard signaling component in the majority of CAR designs in T cells. This served to limit potential confounding effects from this common CAR component. We employed nested GoldenGate assembly to combinatorially generate two comprehensive libraries: 1) a full-length library consisting of 1,332 members each with one to two variable length endodomains, and2) a motif library consisting of 2,756 members each with one to two fixed-length signaling motifs. Both plasmid libraries were used to create lentiviral libraries, which were subsequently transduced into PBMC-derived NK cells at a sufficiently low MOI (MOI = 0.3) to limit integration to a single construct per cell. These libraries provided the basis for the high-throughput screening conducted in this study.

**Figure 1.**
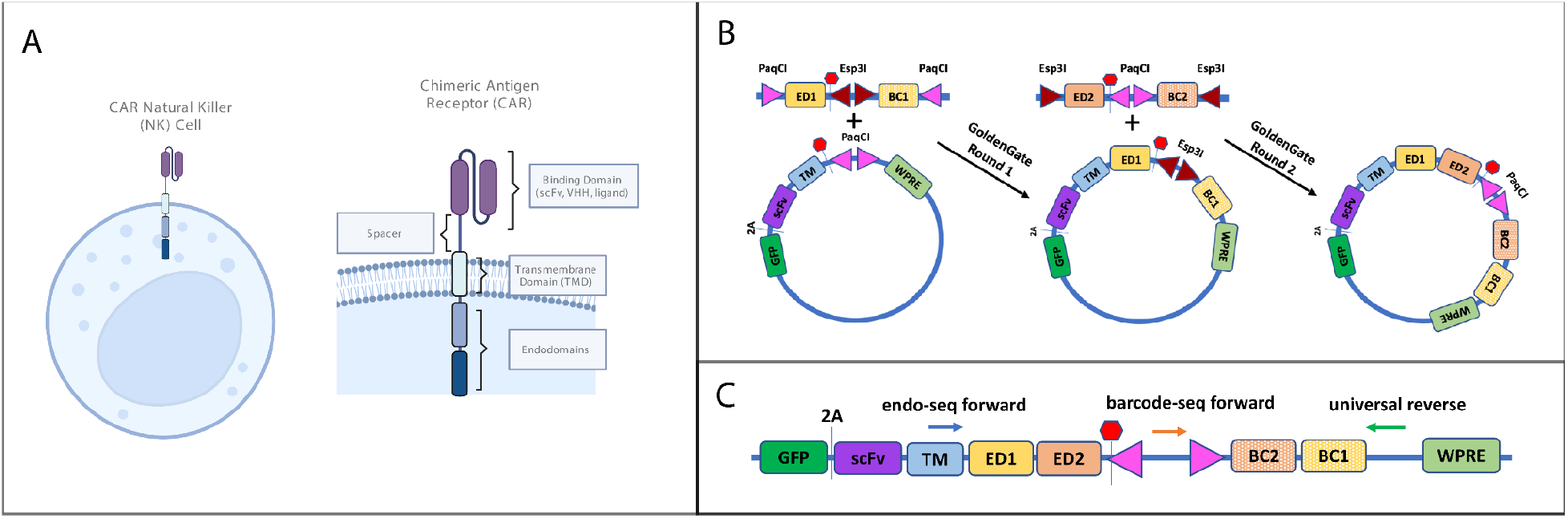
CAR NK design and library construction. a) NK cells were modified to include a chimeric antigen receptor (CAR). CARs comprise an external binding domain, a spacer/hinge, a transmembrane domain, and intracellular signaling domain comprising two endodomains. b) Nested GoldenGate assembly was used to construct our CAR endodomain libraries. In GoldenGate Round 1, gene fragments with endodomain 1 (ED1) and barcode 1 (BC1) were cloned into CAR backbone using PaqCI restriction enzyme and NEBridge Ligase Master Mix. In GoldenGate Round 2, gene fragments with endodomain 2 (ED2) and barcode 2 (BC2) were cloned into the plasmid pool from GoldenGate Round 1 using Esp3I restriction enzyme and NEBridge Ligase Master Mix. c) Sequencing primer positions for endo-seq and barcode-seq.

### CD19-CARs differentially enhanced NK persistence under serial antigen re-challenge

The CD19-CAR full-length and motif libraries were evaluated to assess the capacity of their members to enhance NK cell persistence through a cell-based serial rechallenge assay. The CD19 binder and Raji target cells were selected for their broad applicability to products used in clinical settings to treat hematologic cancers and as a test-bed to enable future binder/antigen combinations. Raji cells were added to cultures at an effector-to-target (E:T) ratio of 1:1 under three protocols: standard media conditions on a daily basis (“Daily Standard”), every two to three days under standard media conditions (“Periodic Standard”), or daily under reduced cytokine support conditions (5 IU/mL, “Daily Low IL-2”). As a control, we maintained cultures in standard media without stimulation with Raji cells in order to rule out antigen-independent proliferation (“Unstimulated”). Our intention was to identify endodomain combinations that could yield superior performance under both standard and low IL-2 conditions. We placed particular emphasis on the identification of endodomains that performed optimally under the Daily Low IL-2 condition, given the increased stringency of this condition and the hypothesis that limited cytokine support is a potential cause of limited *in vivo* persistence. We hypothesized that endodomains that recapitulate IL-2 signaling would drive proliferation and survival under the Daily Low IL-2 condition.

Over an 11-day period, we challenged the pools under the aforementioned conditions (Figure 2A). Following these experiments, the remaining CAR NKs were harvested and sequenced. Utilizing DESeq2^37^, we evaluated changes in the abundance of constructs between antigen re-challenged (treatment) and Unstimulated (control) groups. Our results revealed that serial antigen re-challenges exerted significant selective pressure on CAR-expressing NK cells (Figure 2B and 2C).

**Figure 2.**
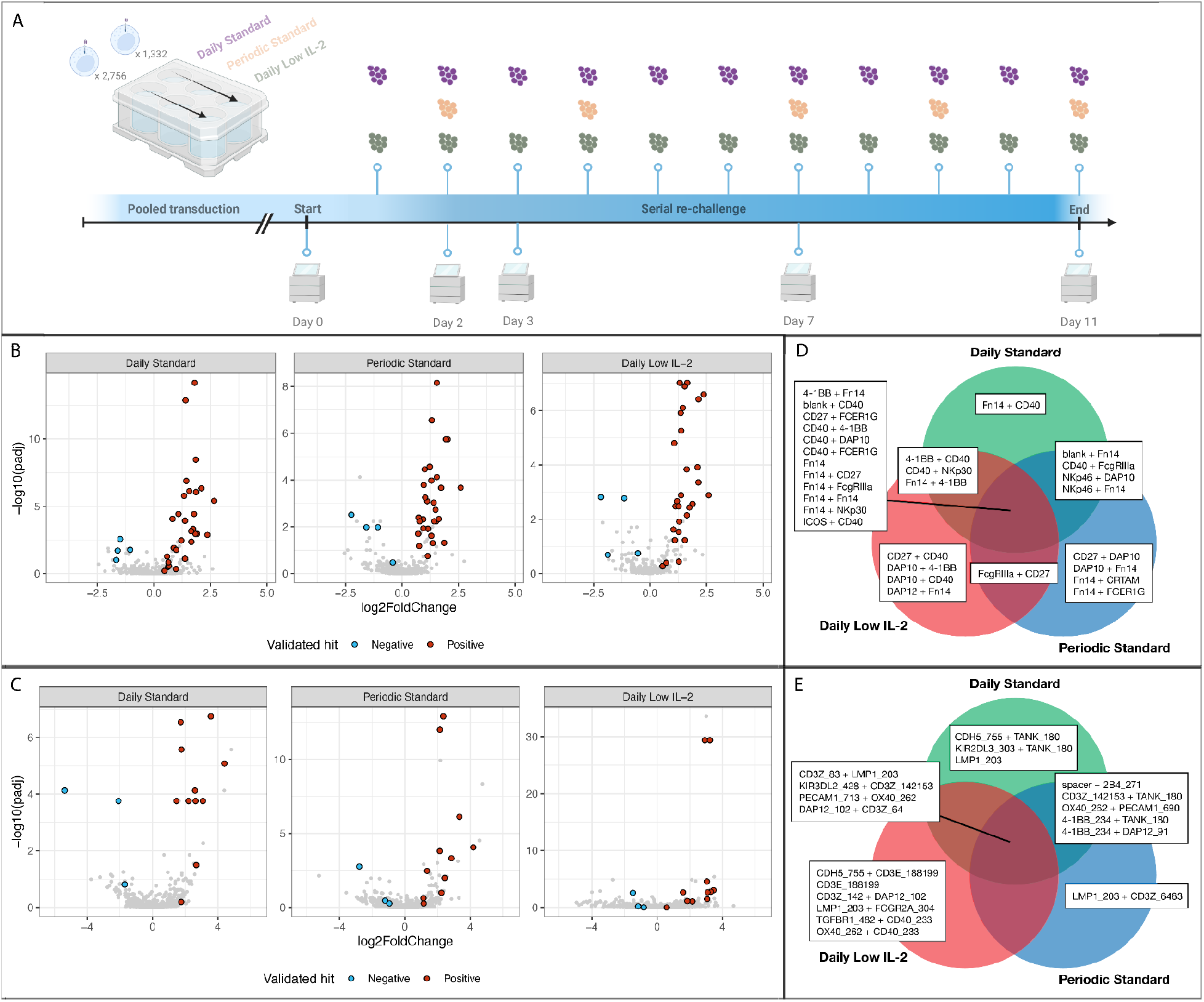
Discovery of CAR NK candidates with pooled screening. a) Screening experiment design. Each library (full-length and motif) was screened in triplicate, twice under normal conditions, with re-challenge occurring at varying intervals, and once in low cytokine conditions. b) Full-length library screen results. The volcano plots show the log2 fold-change and adjusted significance of each construct in the serially stimulated cells compared to control (unstimulated) over the 11-day screen. Positively enriched constructs selected for *in vitro* validation are highlighted in red; negatively enriched constructs are highlighted in blue. c) Motif library screen results. The volcano plots show the log2 fold-change and adjusted significance of each construct in the serially stimulated cells compared to control (unstimulated) over the 11-day screen. d) Top hits from the full-length library. The Venn diagram shows the top 20 constructs which persisted in each screening condition. e) Top hits from the motif library. The Venn diagram shows the top 10 constructs which persisted in each screening condition.

In the full-length library at Day 11, we identified 37 constructs that were significantly enriched (Benjamini-Hochberg adjusted p<0.05) in the Daily Standard condition relative to the Unstimulated condition, and 41 constructs significantly enriched in the Periodic Standard. In the Daily Low IL-2 condition, 33 constructs were significantly enriched. Twelve of the top 20 constructs were shared between all three conditions (Figure 2D). Similar trends were observed for the motif library at Day 11, with 19 constructs significantly enriched under the Daily Standard condition, 30 constructs significantly enriched under the Periodic Standard condition, 25 constructs significantly enriched under Daily Low IL-2 condition, and four of the top 10 were shared between both conditions (Figure 2E). These findings suggest that our library of CARs differentially enhances NK cell persistence in the presence of antigen-expressing cells.

### Barcode-only sequencing reduces length bias in construct recovery

In our profiling of the full-length library, construct lengths appeared to be inversely correlated with read counts. To make sure we captured biologically relevant changes for longer constructs, we separately prepared NGS samples amplifying only the barcodes associated with each endodomain. Barcode-only sequencing recovered more constructs compared to sequencing on both barcode and endodomains. For constructs longer than 200 aa, barcode-only sequencing recovered 549 out of the 659 constructs, compared to 413 constructs recovered with endodomain sequencing (Supplementary Figure 1A). Constructs longer than 200 aa also had higher abundance in barcode-only sequencing than in endodomain sequencing (Supplementary Figure 1B).

### Novel CAR endodomains enhanced persistence of CAR NKs beyond CD3ζ performance

We further aimed to understand the ability of novel CAR endodomains to enhance CAR NK persistence upon antigen re-challenge compared to the conventional CD3ζ endodomain. Using results from barcode-only sequencing to determine the contribution of individual endodomains, we compared the frequency of each endodomain in constructs that were significantly enriched and significantly depleted at the endpoint of the serial re-challenge assay under each growth condition.

In all three assay conditions, endodomains that contained TRAF-binding domains such as Fn14, CD40, CD30, 4-1BB, and CD27, were overrepresented in enriched constructs vs. depleted constructs in the full-length library (Figure 3A). Similarly, in the motif library, TRAF-binding motifs from TANK, OX40, and 4-1BB were overrepresented in the enriched constructs (Figure 3B). In contrast to CAR constructs used for T cells, our CAR backbone did not contain CD3ζ, allowing us to compare the performance of CD3ζ to the other endodomains in CAR NK cells. In the full-length library, Fn14- and CD40-containing constructs displayed higher log fold change enrichment than CD3ζ-containing constructs in the Daily Low IL-2 condition (Figure 3C). In the motif library, TRAF-binding motifs from TANK and OX40 matched the performance of CD3ζ ITAM1 (Figure 3D). Our results show that CD3ζ is not necessary in CAR NK persistence under repeated antigen exposure, underscoring the distinct signaling dynamics between the two cell types. In addition to being over-represented in enriched constructs, high performing constructs displayed a preference for Fn14 at position one (membrane proximal) of the second-generation CAR scaffold. In the Daily Standard condition, 15.6% of enriched constructs had Fn14 at position one vs. 9.4% at position two (cytosol proximal); in the Daily Low IL-2 condition, 17.4% of enriched constructs had Fn14 at position one vs. 10.1% at position two. These observations may have implications for the design and orientation of these endodomains within the CAR construct, suggesting the potential of structural optimization for improved NK cell activation and persistence.

**Figure 3.**
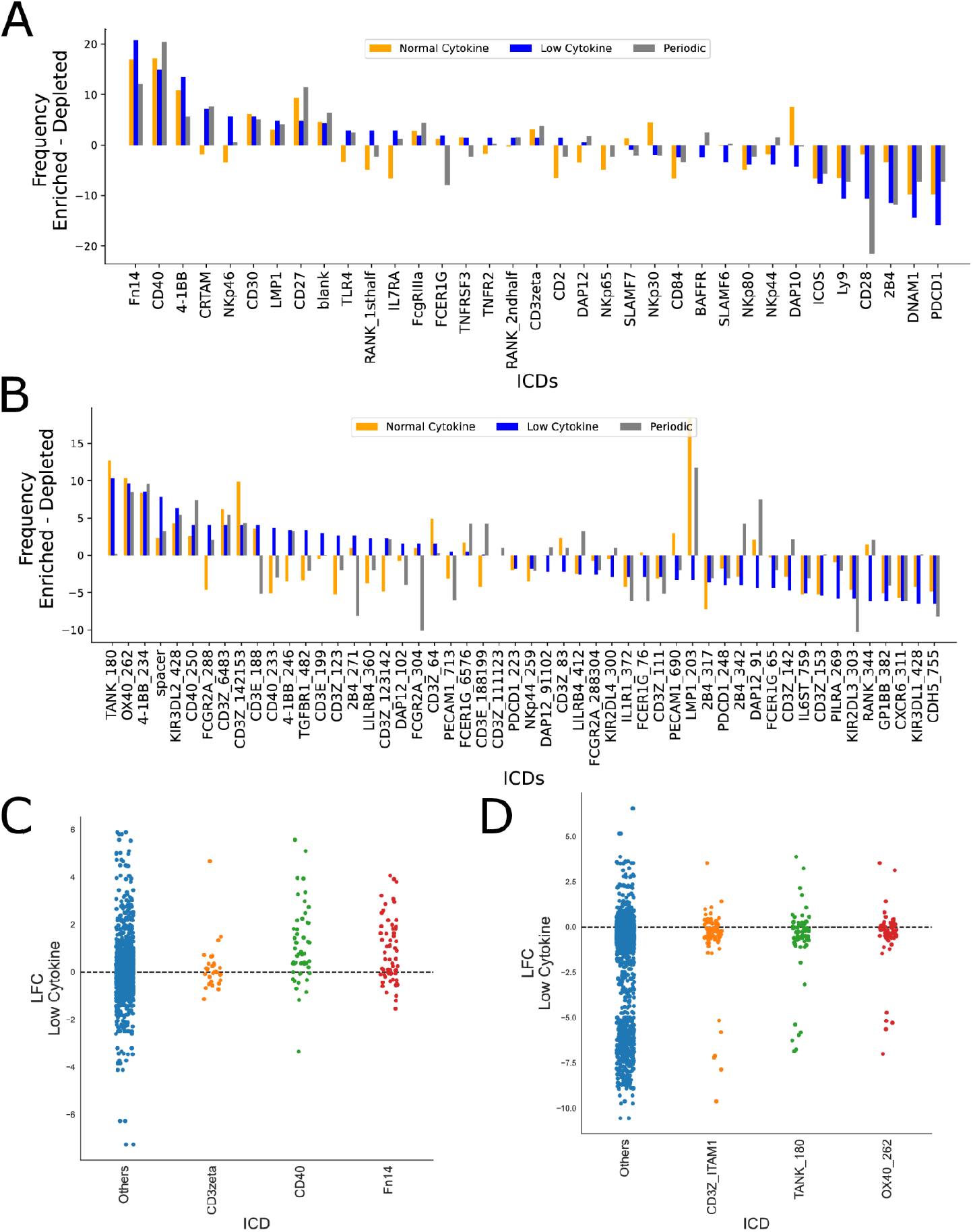
Novel TRAF-binding endodomains outperform CD3ζ. a-b) For each endodomain, the delta between the frequency of that endodomain among significantly enriched constructs vs. significantly depleted constructs was calculated for the full-length library (a) and the motif library (b) under each of the three test conditions (colored bars). c-d) Deseq2 calculated log2 fold changes and BH-adjusted p values (treatment vs. control) at the endpoint of the low cytokine serial antigen rechallenge assay for the full-length library (c) and the motif library (d). All constructs with log2 fold change greater than 0 are plotted, constructs above the horizontal dotted line are significantly enriched. Colored dots highlight constructs that contain Fn14, CD40, and CD3ζ in the full-length library (c) and TRAF-binding motifs in TANK, OX40, and CD3ζ ITAM1,2,3 in the motif library (d).

As a means to better understand the antigen-independent signaling potential of our designs, we also examined enriched constructs in Unstimulated condition at Day 11 versus Day 0. We noted that Fn14 and TANK were not overrepresented, suggesting their enrichment under antigen challenge conditions may indeed be linked to their contribution to antigen-dependent activation and persistence (Supplementary Figure 2).

### Novel CAR endodomains validated for persistence and killing in arrayed format

To validate the enhanced persistence observed in our pooled screen, and to test whether our CD19 CARs additionally increased cytotoxicity, we performed arrayed validation experiments using 40 enriched and seven depleted constructs identified from the screen (Table S3), along with an untransduced (“mock”) control. Lentiviruses expressing each construct were separately prepared to transduce NK cells, and all CARs were expanded following enrichment for CAR expressing cells. Arrayed CAR NKs were rechallenged following the previously described Daily Low IL-2 condition. Survival curves of these CAR-expressing NK cells were determined by flow cytometry-based quantification of Raji/NK cell proportions (Figure 4A).

**Figure 4.**
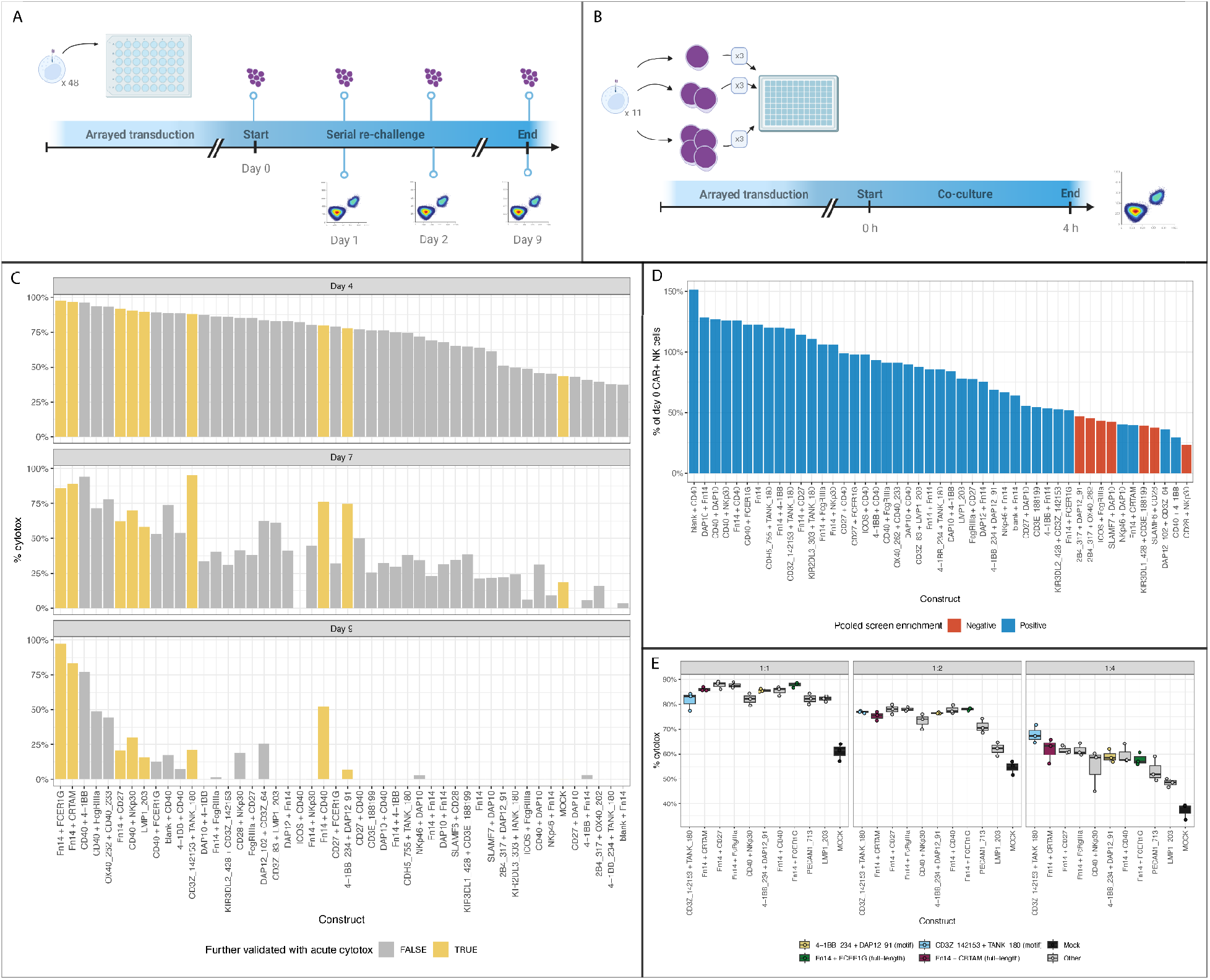
Arrayed *in vitro* validation of screening hits. a) Validation experiment design: 48 CAR NK designs were selected from the full-length and motif libraries (n=29 positive full-length hits, n=4 negative full-length hits, n=11 positive motif hits, n=3 negative motif hits, n=1 mock control). The 48 designs were subjected to a serial stimulation to assay chronic cytotoxicity and intracellular cytokine expression. b) Ten designs were additionally selected for evaluation of acute cytotoxicity at three E:T ratios in triplicate. c) Validation of chronic cytotoxicity. Each bar plot shows the cytotoxicity (measured as the % of Raji cells killed) of each validated design at each of three timepoints during the serial stimulation. Designs highlighted in yellow were selected for further validation of acute cytotoxicity. d) Validation of chronic persistence. Each bar shows the percentage of the initial CAR+ NK cell population remaining at the end of the arrayed serial stimulation assay. Higher bars indicated increased persistence. e) Validation of acute cytotoxicity. The box plot shows the cytotoxicity of each design in each of three E:T ratios after four hours. Highlighted designs were selected for *in vivo* validation.

Analysis of the survival curves demonstrated differential persistence of constructs under re-challenge conditions. Constructs that were enriched in the pooled screen following antigen rechallenge also retained a higher percentage of CAR-positive NK cells at the end of serial antigen rechallenge in the arrayed format (Figure 4D). To ensure that the designs we selected were functionally robust, we also quantified the number of the remaining Raji cells at different timepoints during the antigen re-challenge as a readout of cytotoxicity. Constructs that were enriched in the pooled screen displayed higher cytotoxicity, quantified as percent of dead Raji at Day 4, 7, and 9 of antigen rechallenge (Figure 4C).

In addition to killing in the serial re-challenge assay, we assessed the cytotoxic potential of ten designs in an acute setting (Figure 4B; Table S4). CAR NK cells were incubated with antigen-expressing tumor cells at varied E:T ratios (1:1, 1:2, 1:4) for four hours, keeping the number of target cells consistent across all ratios, while varying the number of effector cells. We observed that constructs enriched in the pooled screen were significantly more effective at killing tumor cells at all E:T ratios than a mock control (Figure 4E; all constructs significant at FDR-corrected p<0.05 in t-tests vs control). In parallel, we confirmed that these designs drove degranulation (CD107a) and IFNɣ secretion in a CAR-dependent manner by intracellular cytokine staining (ICCS; Supplementary Figure 3).

Taken together, these findings validate the potent and enhanced persistence and cytotoxicity of CAR NK cells containing the novel endodomains identified from our high-throughput screen.

### Novel CAR endodomains outperform conventional CARs at killing tumor cells *in vivo*

To evaluate the *in vivo* antitumor efficacy of our identified CAR endodomains, we performed animal experiments using six groups of 12-week-old female NOD.Cg-Prkdcscid Il2rgtm1Wjl Tg(IL15)1Sz/SzJ (NSG-Tg Hu-IL15) mice (four novel CAR endodomains, one ectodomain-only control, and one PBS-only sham control; Table S5), with five mice per group. Each mouse was intravenously administered with 2.5E5 luciferase-expressing Raji cells and then 2.5E6 CAR-expressing NK cells two days later. To monitor the survival and growth of Raji tumor cells, we performed *in vivo* bioluminescence imaging measurements of the average and total luciferase intensity at several time points post NK cell injection (three to five days, and twice more before the end of the study, Figure 5A). Weights were also recorded concurrently. On Day 20 (or earlier if necessary due to tumor burden), mice were euthanized and blood was collected into tubes treated with EDTA/heparin/sodium citrate.

**Figure 5.**
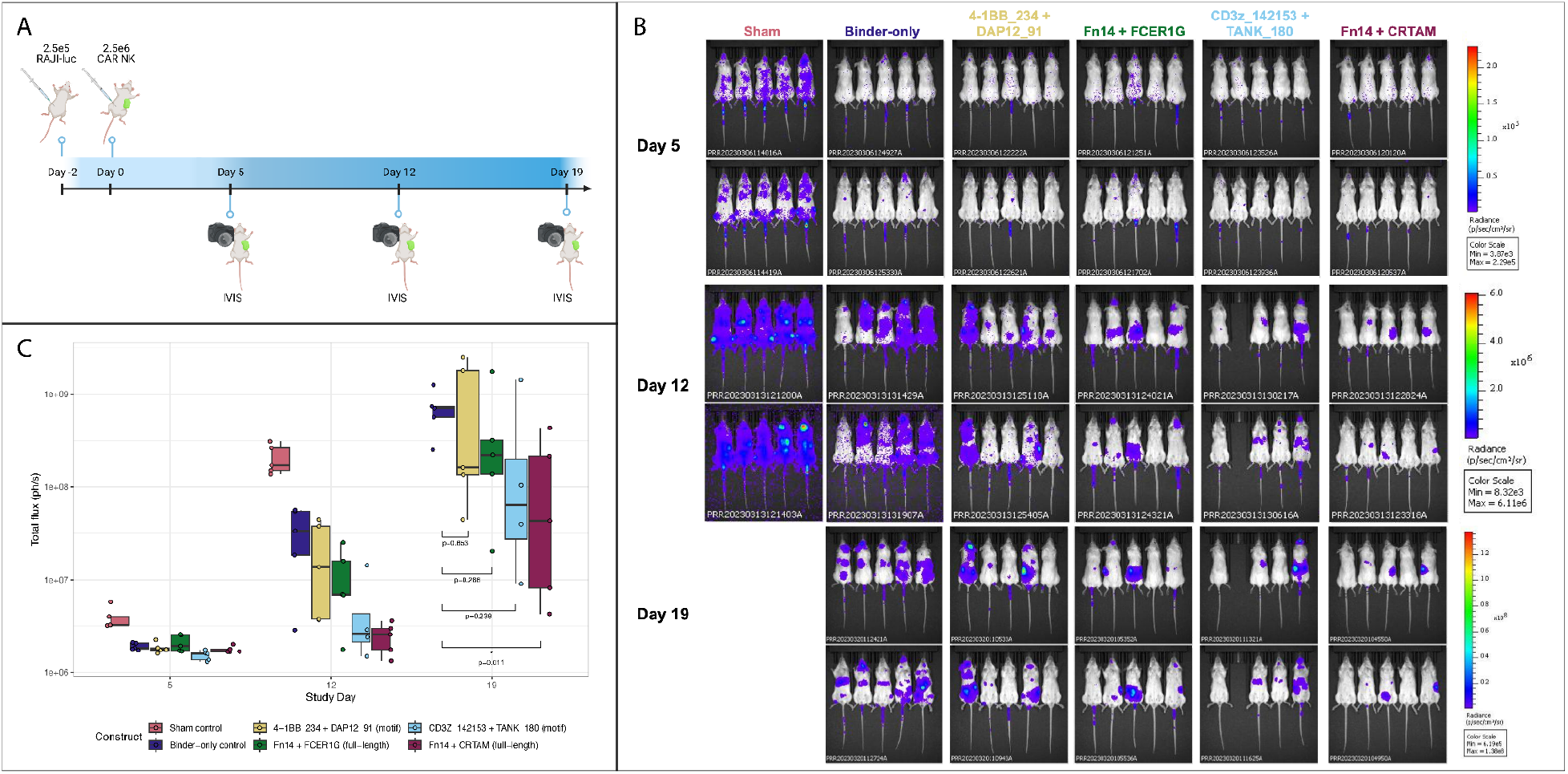
*In vivo* validation of top CAR NK candidates. a) Experiment design: NSG-Tg Hu-IL15 mice were injected with 2.5E5 luciferase-expressing Raji cells, then two days later received a dose of 2.5E6 CAR NK cells. Tumor mass was measured using the IVIS *in vivo* bioluminescence imaging system at three timepoints. b) Bioluminescence imaging. c) Summary of bioluminescence imaging. Box plots show the total flux (anterior and posterior) computed for each mouse (five mice per design and timepoint). Higher values indicate larger tumor mass. The p-values were computed using a t-test between the construct and the binder-only control.

The health status and tumor burden in the mice dictated the course of the experiment. Mice in the sham group had to be sacrificed on Day 15 due to poor health and significant tumor burden, and the mice in the ectodomain-only group were sacrificed on Day 19 for similar reasons. All other mice were maintained until the pre-specified end of the study (Day 20).

Our analysis of *in vivo* imaging data showed a notable difference in tumor burden as measured by luciferase intensity across different groups. Specifically, mice receiving CAR NK cells expressing either Fn14+CRTAM or CD3ζ ITAM3+TANK TRAF binding domain endodomains showed lower tumor-associated luciferase intensity at Days 5, 12, and 19 post NK cell injection, compared to the other groups (Figure 5B-C). This data suggests that the inclusion of novel CAR endodomains can significantly enhance *in viv*o antitumor efficacy of NK cells.

## Discussion

Here, we present an innovative approach for unbiased screening of diverse libraries to uncover novel CAR NK functionality. Using a pooled screening approach, we identified several novel CAR endodomains that significantly enhanced NK cell persistence and cytotoxicity in low cytokine-support conditions, most notably two TRAF (TNF receptor-associated factor) binding proteins: Fn14 and TANK. These findings provide new avenues for augmenting the efficacy of CAR NK cell therapies.

While we identified some constructs that persist only under low cytokine support, there was a considerable amount of overlap in construct performance across conditions (Figure 2C, 2E). This may be due to the converging signaling pathways downstream of the endodomains in the current library. Future studies featuring libraries with greater diversity in terms of signaling domains are required to decouple the effects of signaling on persistence with or without IL-2 support.

TANK emerged as a significant novel hit in our investigation, suggesting that TANK potentially bolsters the persistence and cytotoxicity of NK cells. The protein encoded by TANK is known to interact with the tumor necrosis factor receptor-associated factor (TRAF) family of proteins^7, 38, 39^. This interaction mainly takes place in the cytoplasm and can involve TRAF1, TRAF2, or TRAF3. By binding to these TRAF proteins, TANK effectively inhibits their function, sequestering them into a latent state within the cytoplasm. An extensive mechanistic exploration is warranted to clarify these inferences, and the role of this molecular mechanism within the context of enhancing NK cell function presents an exciting avenue for future investigation.

Fn14, another TRAF-binding protein in our library, is known to play an important role in cell proliferation, an essential characteristic for the effective functioning of therapeutic NK cells^40–42^. By enhancing proliferation, Fn14 may help to augment the cytotoxic capabilities of CAR NK cells, rendering them more effective against cancer cells.

Unlike T cells, CAR NK cells do not rely on CD3ζ for activation. A similar pooled endodomain screen in CAR T cells yielded CD3ζ as one of the top endodomains for T cell activation (unpublished data). Together, these findings are consistent with the hypothesis that T cells and NK cells are activated by different signaling cascades, and optimizing CARs in an NK context yields more functional designs than CARs discovered in a T cell context. Given the distinct signaling dynamics of NK cells, CARs discovered in an NK context, when used in CAR NKs, outperform CARs discovered in a T cell context. Our future research will focus on screening combinations of endo-, ecto-, and transmembrane domains to further enhance the efficacy of CAR NK cell therapies.

## Materials and Methods

### Design of full length and motif libraries

We compiled a list of endodomains from literature and genome mining and narrowed down the list to 36 full-length endodomains and 52 signaling motifs, derived from human endodomains. The 36 full-length endodomains were compiled from extensive literature search of CAR T, NK cell, NFAT, AP1, and NF-kB signaling publications.

To construct the fixed-length motif library, we focused on signaling proteins containing the Src Homology 2/3 (SH2/SH3) domain and gathered 41 immunoreceptor tyrosine-based regulatory motifs, including 14 inhibitory (ITIM), 22 activating (ITAM), and five switching (ITSM) motifs^43^. Additionally, we identified 10 TRAF-binding domains from various receptors and intracellular proteins that interact with TRAF 1, 2, 3, 5, and 6^32, 44, 45^. Because ITAMs naturally occur in tandem with the consensus sequence YxxI/Lx(6–12)YxxI/L, we preserve this pattern for the known ITAMs in our library and also repeated all the other motifs two times in tandem to arrive at a fixed length (36AA) for the entire library.

### GoldenGate assembly of libraries

We used nested GoldenGate assembly to combinatorially generate two libraries: 1) a 1,332 member, variable-length full-length endodomain library and 2) a 2,756 member, fixed-length signaling motif endodomain library. Two rounds of GoldenGate assembly were essential for incorporating a unique DNA barcode for each endodomain. DNA barcodes are intended to be used in Illumina next generation sequencing (NGS) to minimize the length bias in PCR and Illumina flowcell hybridization. First, we assigned each endodomain with a unique DNA barcode. Second, we added Esp3I GoldenGate cloning sites between each endodomain and its barcode and added PaqCI GoldenGate cloning sites flanking on both ends (Figure 1b). Third, we ordered these GoldenGate gene fragments as eBlocks from IDT. For the first round of GoldenGate assembly, we pooled all the eBlocks at equal molar ratio and used PaqCI Type IIs restriction enzyme (NEB), NEBridge Ligase Master Mix (NEB), and the cloning backbone containing PaqCI sites (Figure 1b). GoldenGate reaction was run according to NEBridge protocol and transformed into NEB10β cells (NEB), and recovered in 50mL of TB media with Carbenicillin overnight at 37°C shaking incubator. Plasmid pool from bacteria culture was maxiprepped using GeneJet endofree maxiprep kit. For the second round of Goldengate, we ordered a second set of eBlocks with Esp3I as flanking GoldenGate cloning sites and PaqCI as internal GoldenGate cloning sites (Figure 1b). We pooled the second set of eBlocks at equal molar ratio and used Esp3I Type IIs restriction enzyme (NEB), NEBridge Ligase Master Mix, and the maxiprepped plasmid pool from the first round of GoldenGate assembly as cloning backbone. We repeated the same GoldenGate reaction, transformation, bacteria culturing, and plasmid prep steps from the first round for the second round. The second round GoldenGate assembled plasmid pool was sequenced by NGS to determine the distribution of all expected plasmids in the pool.

### Cell lines

Lenti-X 293T (human embryonic kidney) cells were purchased from Takara Bio Inc, and cultured in Dulbecco’s Modified Eagle’s Medium (DMEM; ThermoFisher Scientific Waltham, MA), and supplemented with 10% fetal bovine serum (FBS, HyClone, Logan, UT), 1% Minimum Essential Medium Non-Essential Amino Acids (MEM-NEAA, Gibco), 1% Sodium Pyruvate (NaPyr, Gibco), 1% Glutamax (Gibco).

Raji (Burkitt’s lymphoma) cell lines were purchased from the American Type Culture Collection (ATCC, Manassas, VA) and cultured in Roswell Park Memorial Institute (RPMI 1640; ThermoFisher Scientific) and supplemented with 10% FBS, 1% MEM-NEAA, 1% NaPyr, and 1% Glutamax.

Rajis used for co-culture assays were transduced with a lentiviral vector carrying LSSmOrange and mThy1.2 tag. LSSmOrange cells were isolated with mThy1.2-APC antibody (BioLegend) and EasySep APC Positive Selection Kit II (StemCell Technologies) following manufacturers recommendations.

All cells were cultured in a humidified atmosphere containing 5% CO2 at 37°C.

### Lentivirus generation

The day before transfection, Lenti-X 293T cells were seeded at a density of 1.2E7 cells per T182 TC treated flask. Transfections were carried out using the TransIT-293 reagent (Mirus Bio) following the manufacturer’s instructions. Plasmid DNA was mixed in a molar ratio of 4:2:1:1 for the transfer plasmid, GagPol RRE, Rev, and pMD2.G plasmids respectively. A DNA to lipid ratio of 3:1 was used, equating to 6 µL of TransIT reagent for every 2 µg of plasmid DNA.

Three days post-transfection, the viral supernatant was harvested and centrifuged at 1500 x g for 10 minutes. The supernatant was then cleared further by passage through a 0.45 µm PES filter. The viral particles in the supernatant were then concentrated using the LentiX Concentrator (Takara Bio), in accordance with the manufacturer’s instructions.

### Primary NK cell isolation

Human NK cells were isolated directly from a fresh leukopak by CD56-based, positive immunomagnetic enrichment (STEMCELL #17855, EasySep™ Human CD56 Positive Selection Kit II) and were frozen down in CS10 CryoStor cell cryopreservation media media (Sigma-Aldrich).

### Primary NK cell expansion

NK cells were cultured in CTS NK-Xpander Complete Culture Media (Gibco) supplemented with 5% human serum (Sigma-Aldrich) and 50 U/mL Interleukin-2 (IL-2, R&D Systems). Pre-isolated NK cells were thawed and co-cultured at a 1:1 effector:target (E:T) ratio with mitomycin C-treated K562 cells that were genetically engineered to express membrane-bound Interleukin-21 (mbIL-21) and 4-1BBL. The culture medium was refreshed every three to four days, and the cells were cocultured with mitomycin C-treated cells on a weekly basis.

### Primary NK cell transduction

Cells were transduced 10 days post thaw. Prior to transduction, 4E7 cells were resuspended in 4 mL NK cell expansion media with 3 uM BX-795 (Sigma-Aldrich), and were left to incubate in a 37°C incubator for 30 minutes. Following incubation, 2mL of NK cell expansion media containing 40ug/mL of polybrene (Sigma-Aldrich) was added to cells. The cells were distributed across TC treated 96-well u-bottom plate (150uL/well) and 50uL of pre-diluted virus in NK cell expansion media was added to each well. The 96 well plate was spinoculated at 1200xg for 30 minutes at 32°C in a centrifuge. After the spinoculation, the cells were mixed and incubated in a 37°C incubator for one hour. Post incubation, we performed media exchange, and recovered the cells in a six-well GREX plate (Wilson Wolf, New Brighton Minnesota, USA).

### CAR NK cell enrichment

We added a FLAG-tag at the N-terminus of our CAR for detection and magnetic enrichment. On Day 14, three days post-transduction, we performed CAR NK cell enrichment using FLAG-APC antibody (BioLegend) and EasySep APC Positive Selection Kit II (StemCell Technologies) following manufacturers recommendations.

### Flow cytometry

NK cells were identified via flow cytometer using CD56-APC antibody. Transduction efficiency was estimated using GFP marker in the CAR plasmid. CAR expression was estimated using FLAG-APC antibody. Rajis were estimated by LSSmOrange marker. Human Fc receptors were blocked using Human TruStain FcX (Biolegend).

| Marker | Fluorophore | Clone | Company |
| --- | --- | --- | --- |
| FLAG | APC | L5 | Biolegend |
| CD56 | APC | NCAM | Biolegend |
| IFNγ | PE | 4S.B3 | Biolegend |
| CD107a | PE-Cy7 | H4A3 | Biolegend |

### Serial restimulation assay

For pooled co-culturing assay, CAR NK cells were cultured in four different assay conditions: Daily Standard, Periodic Standard, Unstimulated, and Daily Low IL-2. In the Daily Low IL-2 condition, 5 IU/mL of IL-2 was used, while 50 IU/mL of IL-2 was used in the other three conditions. We stimulated CAR NK cells with Rajis at 1:1 E:T ratio in all conditions except in the Unstimulated condition, in which there were no Raji. For the Daily Standard and Daily Low IL-2 conditions, Rajis were added daily. For the Periodic Standard condition, Rajis were added every two to three days. The pooled coculture assay was done in 24-well G-REX plates in triplicates. We started with 1E6 CAR NK cells and 1E6 Rajis in every well for the three stim conditions and added Rajis as needed to reset the E:T ratio back to 1:1 over 11 days of serial restimulation assay. For the Unstimulated condition, we simply added fresh media every two to three days. CAR NK-cell proliferation and cytotoxicity were measured using flow cytometric analysis. CAR NKs were collected daily for genomic DNA (gDNA) isolation, followed by amplicon sequencing to measure CAR enrichment over time.

For arrayed validation, serial cytotoxicity and persistence were validated in an arrayed format using an otherwise identical serial restimulation assay. To validate which endodomains could recapitulate IL-2 signaling, we rechallenged cells under the previously described Daily Low IL-2 condition. For the Unstimulated condition, the setup was nearly identical to the schema used in the pooled setting, and only differed in terms of the amount of cytokine supplemented (5 IU/mL as opposed to 50 IU/mL.Target cells only condition was also run alongside to inform Raji growth kinetics in coculturing assay. CAR NK cell expansion and cytotoxicity were estimated using flow based measurements. Cytotoxicity was estimated by (Raji(est) - Raji(actual)/Raji(est))*100, where Raji(est) was derived from the target cells only condition.

### Acute cytotoxicity assay

CAR NK cells were cocultured with Rajis at indicated E:T ratios and target only conditions for four hours. CAR NK and Raji estimates were measured via flow cytometric analysis. Cytotoxicity was estimated using following formula: %Lysis = ((Rajis(target only) - Rajis(+CAR))/Rajis(target only))*100.

### Acute cytokine secretion assay

To assess the release of cytokines and degranulation by CAR-expressing NK cells, the cells were first co-incubated with target cells. During this four-hour incubation, the cells were also exposed to a cocktail of CD107a-PECy7 (Biolegend), 1X Monensin (eBioscience), and 1X BFA (eBioscience). Subsequently, the cells were stained with CD56-APC (Biolegend) for 20 minutes at room temperature (RT). Following the surface staining, the cells were fixed and permeabilized using Cyto-Fast™ (Biolegend) Fix Buffer for 20 minutes at RT. Thereafter, intracellular staining for IFNɣ-PE (Biolegend) was performed to identify cytokine-producing cells. The staining process provides a snapshot of the cytokine secretion by the CAR-expressing NK cells following the interaction with the target cells.

### Illumina sequencing

Cells were lysed in Qiagen’s RLT + DTT buffer. gDNA was harvested from lysate using Quick-DNA Micro Plus Kit (Zymo Research) following the manufacturer’s instructions. A two-step PCR was performed on the isolated gDNA. For preparation of endodomain sequencing libraries, PCR1 primers flanked the endodomain and barcode region of the plasmid. We used NEB Q5 polymerase for 25 cycles followed by 1.0X SPRI purification. PCR2 primers included P5 and P7 Illumina sequencing adaptors and sample-specific barcodes. For PCR2, we used NEB Q5 polymerase for six cycles followed by 1.0X SPRI purification. Illumina libraries were normalized and pooled and sequenced on an Illumina iSeq with a 2X150bp reads run mode. For barcode-seq prep, we used PCR1 primers that flanked the barcode region, conducted 2.0X SPRI purification for small amplicons in both PCR1 and PCR2, and used 2×75bp reads run mode on the Illumina iSeq.

### Processing and analysis of NGS data

Reads were trimmed of primers, enzymatic restriction sites, and flanking GoldenGate linkers using Cutadapt v4.1^46^.

In libraries generated with our endo-seq prep, after trimming, R1 contained sequence from one or both endodomains, and R2 contained sequence from the second endodomain. Reads were aligned to our reference using BWA^47^. We counted the number of reads in which we observed each alignment or pair of alignments to quantify construct abundance in each sample.

In libraries generated with our barcode-seq prep, trimmed reads were merged using PEAR^48^. The remaining sequence contained one or two barcodes interspersed with a fixed linker sequence. Trimmed and merged reads were decomposed into the constituent barcodes, which were then matched to our reference, permitting an edit distance of one. We counted the number of reads in which we observed each barcode or pair of barcodes to quantify construct abundance in each sample.

To quantify differential expression of constructs with and without serial antigen re-challenge, we applied DESeq2^37^ to the construct count matrix. Specifically, construct counts from the endpoint of the serial antigen re-challenge screen were collected from three replicates in each of four conditions (Daily Low IL-2, Daily Standard, Periodic Standard, Unstimulated). The likelihood ratio test was used to estimate the log fold change and the false discovery rate-adjusted p-value for each construct between each treatment condition and the control condition (no tumor cell) over the course of the time series.

### Xenograft model

50 female, 12 week-old NOD.Cg-Prkdcscid Il2rgtm1Wjl Tg(IL15)1Sz/SzJ (Strain #:030890) (NSG-Tg Hu-IL15) mice were obtained from Jackson Laboratory. Mice were injected with 2.5E5 luciferase-expressing Raji cells two days prior to CAR NK injection (Figure 4). On Day 0, 2.5E6 CAR NK cells were injected into all but the sham control group, which received an injection of PBS vehicle only. Tumor mass was measured using the IVIS *in vivo* bioluminescence imaging system at three timepoints: Day 5, Day 12, and Day 19. Weights were also recorded concurrently. On Day 20 (or earlier if necessary due to tumor burden), mice were euthanized and blood was collected into tubes treated with EDTA/heparin/sodium citrate.

## Acknowledgments

This research was supported by the Genomics & Bioinformatics Shared Resource (RRID:SCR_022606) of the Fred Hutch/University of Washington/Seattle Children’s Cancer Consortium (P30 CA015704). Research reported in this publication utilized the Preclinical Research Shared Resource at Huntsman Cancer Institute at the University of Utah and was supported by the National Cancer Institute of the National Institutes of Health under Award Number P30CA042014. The content is solely the responsibility of the authors and does not necessarily represent the official views of the NIH. Figures 2a, 3a, and 4a were created with BioRender.com.

## Author Contributions

M.W., A.C., M.Y., B.B., L.G., B.D., and M.D. designed the study. A.C. designed CAR plasmids. M.W., A.C., N.K., R.T., N.B., and M.Y. performed material generation, sequencing, screening, and validation. M.S. and L.G. performed computational analysis. M.W., A.C., M.S., L.G., B.D., and M.D. prepared the manuscript.

