## Supplemental Figures for "TRAFfic signals: High-throughput CAR discovery in NK cells reveals novel TRAF-binding endodomains that drive enhanced persistence and cytotoxicity"

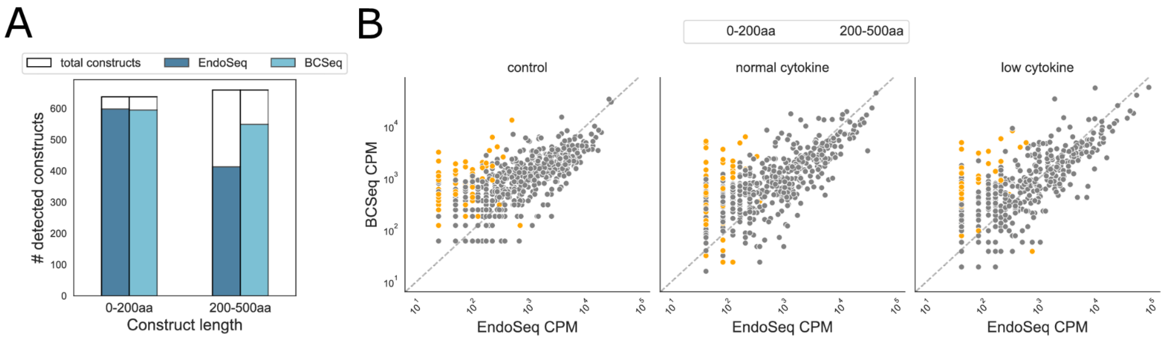


**Supplementary Figure 1. Barcode-only sequencing enhances recovery of longer constructs.** a) Barcode-only sequencing recovered 1,144 whereas endo+barcode sequencing recovered 1,011 out of 1,296 total GenII constructs. b) Read abundance (counts per million) for each construct in endo+barcode sequencing vs. barcode-only sequencing. Constructs longer than 200aa (orange) had higher abundance in barcode-only sequencing.

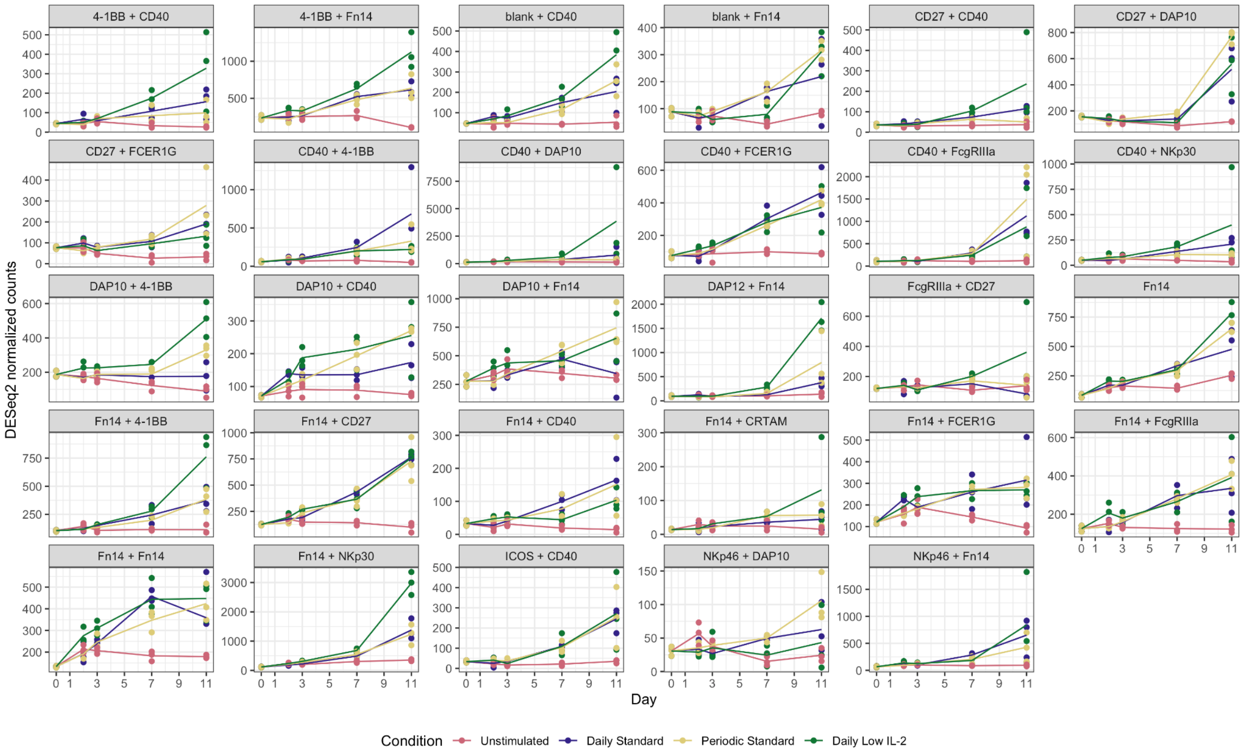


**Supplementary Figure 2. Evidence of antigen-dependent signaling.** Each facet shows normalized abundance over the course of the pooled screen time course of the positive hits selected for arrayed validation.

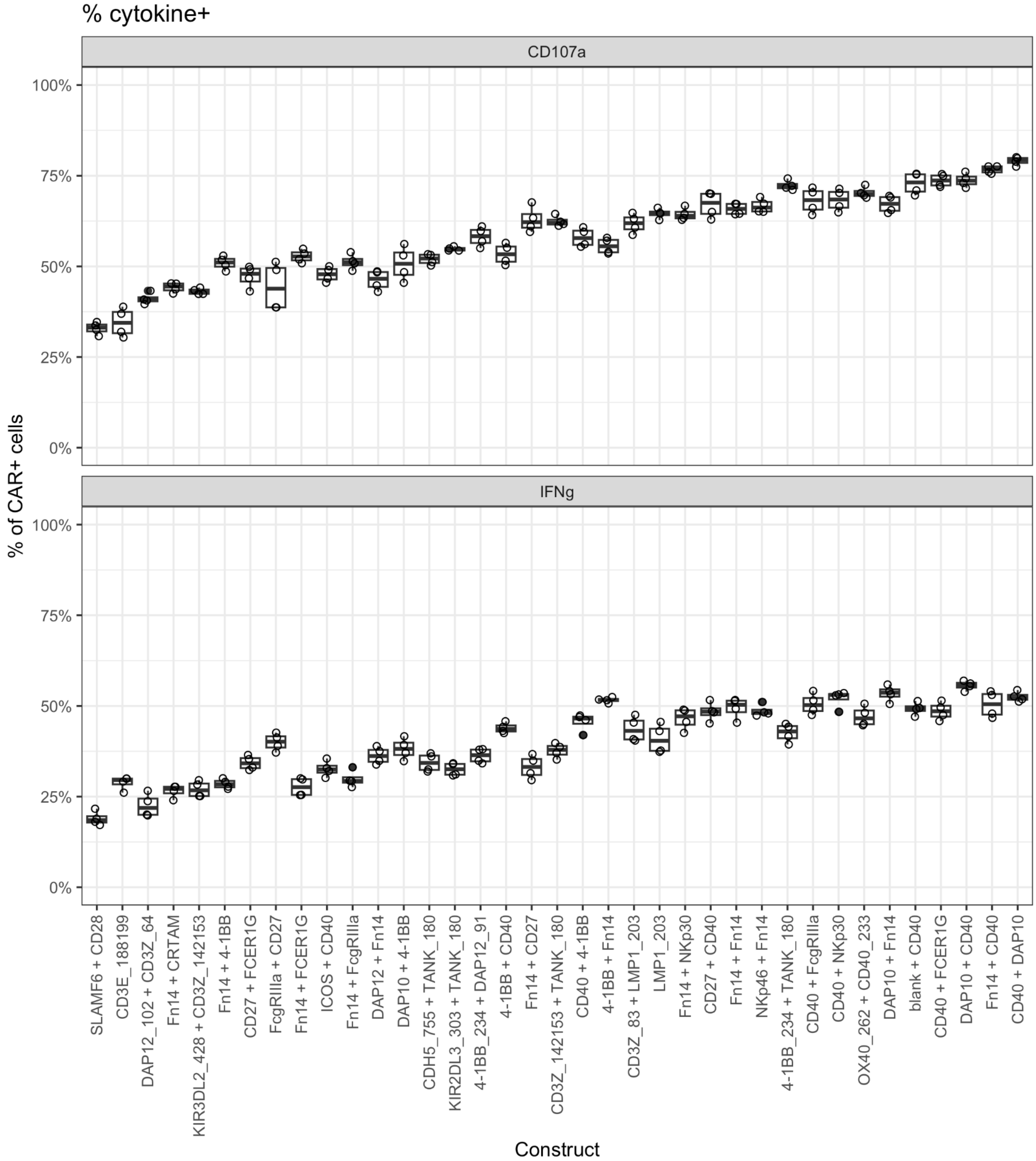


**Supplementary Figure 3. Intracellular cytokine staining of validated constructs.** a) % of CAR+ cells expressing CD107a. b) % of CAR+ cells expressing IFNɣ.
